# Comparisons of paternity success for resident and non-resident males and their influences on paternal sibling cohorts in Japanese macaques on Shodoshima Island

**DOI:** 10.1101/2023.02.14.528291

**Authors:** Shintaro Ishizuka, Eiji Inoue

## Abstract

In group-living mammals, reproductive success can be attributed to both resident and non-resident males. However, the reproductive success of non-resident males has rarely been investigated at an individual level. As male reproductive success is highly skewed towards specific males, often the most dominant males, the percentage of similar-aged paternal siblings within groups is expected to increase. However, the extent to which each male contributes to the production of cohorts of paternal siblings remains unclear. Here we examined the paternity of 46 offspring born over five consecutive years in a group of Japanese macaques *Macaca fuscata* on Shodoshima Island. We quantitatively assessed paternity success for males, including both resident and non-resident males, and the percentages of paternal sibling dyads in the same age cohorts produced by those males. Non-resident males sired neither higher nor lower percentage of offspring compared to resident males, indicating that various males, including non-resident males, usually partake in the within-group breeding of macaque groups. These are possibly because female preference of mating partners can change over time. Subadult males had a lower percentage of paternity success, which may be because females may not prefer the physically immature subadult males. Various males, including non-resident males, contributed to the creation of paternal sibling in the same age cohort. The overall results suggest that not only resident but also non-resident males play an important role in shaping within-group kin structures. Future studies are required to examine how paternal siblings interact with each other.

## Introduction

Males of group-living mammals often achieve different degrees of reproductive success within a group (Cowlishaw and Dunbar 1991; Kutsukake and Nunn 2009). Although resident males tend to attain large amounts of reproductive successes in many species, males from outside the group also find reproductive success within the group. The paternity of offspring assigned to males outside a group is termed extra-group paternity (EGP). EGP is a phenomenon widely observed in various mammalian species (e.g. primates: Ruiz-Lambides et al. 2017; Miller et al. 2021; carnivores: Lyke et al. 2013; Nicholas et al. 2015; rodents: Wells et al. 2021; bats: Ortega et al. 2003; shrews: 152. South et al. 2007) and is essential for the evolution of social systems in animals as it increases genetic diversity within groups and is consequently associated with the increased chance of survivorship (Reed and Frankham 2003). EGP has been usually determined when the paternity of offspring is unsuccessfully assigned to all candidate fathers within a group, probably because individual identification and sample collection for non-resident males are usually difficult in the field. Thus, the overall percentages of EGP have been well investigated (Isvaran and Clutton-Brock 2007; Miller et al. 2021). However, EGP has rarely been assigned to a certain non-resident male, so paternity success for each non-resident male has not been quantitatively assessed at the individual level. Thus, the extent to which each non-resident male has a probability of siring offspring and how that may change remains unclear. To better understand male reproductive strategies with reference to the social statuses in group-living mammals, it is necessary to assess the reproductive success for not only the resident males, but also the non-resident males, by individually identifying and sampling both these types of males.

Variance in the reproductive success of males modulates the abundance of paternal kin in animal groups. In mammalian species, kinship is well known to modulate social relationships among individuals because kin selection favors reciprocity and cooperation with kin over non-kin (Hamilton 1964; Silk 2009; Smith 2014). Although it is unclear whether individuals can discriminate paternal kin in promiscuous species, evidence of paternal kin bias behavior has been reported across various mammalian species (e.g. primates: Smith et al. 2003; Cords et al. 2018; De Moor et al. 2020; carnivores: Wahaj et al. 2004). Paternal kin bias is often based on familiarity among similar-aged individuals (Widdig 2007). In animal groups, the percentage of paternal sibling dyads in the same age cohort increases in hand with the male reproductive skew, and breeding males are frequently replaced (Altmann 1979; Lukas et al. 2005; Widdig 2013). Therefore, age proximity has been shown to be utilized for paternal kin discrimination (Widdig et al. 2001). Quantitative assessments of the percentage of paternal sibling dyads in the same age cohorts are therefore important for the interpretation of paternal kin bias in promiscuous mammals. To date, most studies have focused on the contributions of the most dominant males to the percentages of paternal sibling dyads in the same age cohorts (Langergraber et al. 2007; Widdig 2013; De Moor et al. 2020), since they were presumed to attain the largest amounts of reproductive success, thereby neglecting the contributions of other males. Therefore, for a comprehensive understanding of the mechanisms that produce kin-dyads within groups of social animals, examining the percentages of paternal sibling dyads produced by resident males beside the most dominant males and non-resident males at an individual level is crucial.

The Japanese macaques *Macaca fuscata* are an ideal species to fill this knowledge gap. They form female-philopatric and multimale/multifemale groups (Yamagiwa and Hill 1998). First, we can assess the likelihood of siring for each non-resident male. During the mating season, several non-resident males sporadically appear around the group and copulate with the group’s females (Takahashi 2001; Hayakawa 2008). Paternity results have confirmed that such copulations can result in EGP (Soltis et al. 2001; Hayakawa 2008). Considering that previous field research succeeded in collecting behavioral data and genetic samples from non-resident males (Inoue and Takenaka 2008; Hayakawa 2008; Kawazoe 2016), it may be possible to expand on it further to examine the paternity success for non-resident males at the individual level. Second, previous studies have shown that the percentage of the alpha male paternity is usually low (Inoue et al. 1990; Inoue and Takenaka 2008; Hayakawa 2008), although one study reported a relatively high percentage (Soltis et al. 2001). This suggests that both alpha and non-alpha males produce paternal sibling dyads and that the percentages of paternal sibling dyads produced by either type of male can be assessed quantitatively. In this study, we examined the paternity of the offspring born in five consecutive years within a group of Japanese macaques, wherein both resident and non-resident males were identified and sampled. We assessed the percentages of paternity success for both resident and non-resident males and the percentages of paternal sibling dyads produced by them among the offspring in each year.

## Materials and Methods

### Study subjects

The study subjects were free-ranging Japanese macaques of the B-group inhabiting an area near the Choshikei Monkey Park on Shodoshima Island (Ishizuka 2021). The macaques are provisioned by park staff at approximately 8:30 h, 14:00 h, and 16:00 h and by tourists visiting the park. We have conducted continuous field research on this group since 2017. Most group members, including all the adult males, adult females, and their offspring, were fully identified in 2019, and their demography has been continuously recorded since February 2019. The group comprised approximately two hundred individuals in 2017, after which the number of group members gradually decreased until 2020, probably because several members were captured for wildlife management purposes. In 2022, the group consisted of sixty-eight individuals.

### Classification of males

This study focused on males aged ≧4 years assumed to be sexually mature (Takahata et al. 1998) and were classified based on their age class and social status. Following a previous study (Inoue and Takenaka 2008), the age class of males aged 4–6 years and that > 6 years was categorized as “subadult” and “adult,” respectively. For this study, the males were first divided into resident or non-resident males, based on their social status. As a part of their daily monitoring, the park staff and SI recorded the presence and movement of each male in the group. When certain males were observed throughout a year, they were classified as resident males. Conversely, when certain males were observed only during mating seasons, they were classified as non-resident males. The social status of resident males was further divided into dominant or subordinate males. As Hill (1999) reported, males of provisioned groups are constantly being brought into competition with one another over food, which emphasizes the dominance-subordinate aspects of their relationships. In our study, some males were observed to access to preferred food at the provisioning area, whereas other males were driven away by both the males and females who stayed at the provisioning area and were thus rarely observed to enter into the area. Therefore, we classified those males that were allowed to access the provisioning area as dominant and those that were driven away from the provisioning area as subordinate males. Collectively, the social statuses of males were classified into three categories: dominant, subordinate, and non-resident males. This is in accordance with previous studies on macaque species (e.g. Hayakawa 2008; Toyoda et al. 2022). Furthermore, the alpha male, which is the most dominant male, was identified each year as one among the dominant males. The alpha male in groups of Japanese macaques can be identified by their status-typical behaviors, such as support of losers rather than winners at intra-group competitions (Watanabe 1979) or taking the lead in attacking members of different groups (Maruhashi 1982). For this study, the alpha males were identified to assess the percentage of their paternity and its contribution to the production of paternal siblings. Information for candidate fathers is shown in Table S1.

Notably, the status of the alpha male changed drastically during the 2017 mating season. The alpha male at the beginning of the mating season was “TR,” who became the alpha male in December 2015. However, he got injured in a serious attack by “SB,” who was the second-ranking male at the time, and lost the position of alpha on the 13th of October 2017. “SB” was considered as the alpha male successor until he suddenly disappeared from the troop seven days after the altercation. Thus, another male called “SN” assumed the position of alpha male from the 20th of October 2017. Due to these complex changes in the alpha male status, we did not categorize any single male as the alpha in the 2017 mating season. A few years later, in June 2021, “SN” suddenly disappeared from the troop and “TY” became the next alpha male.

### Genetic analysis

Feces, saliva, and sperm samples were collected using cotton swabs and stored in lysis buffer at ambient temperature for genetic analysis. DNA was extracted from the samples using a QIAamp Stool Mini Fast Kit (Qiagen, CA, USA). Using DNA extracts, the genotypes at 16 autosomal microsatellite loci were analyzed. Multiplex PCRs were performed for genotyping (Arandjelovic et al. 2009), using three primer sets: set A (D3S1768, D5S820, D6S501, D17S1290, MFGT18, and MFGT22), set B (D6S493, D314S306, D19S582, MFGT5, and MFGT21), and set C (D1S548, D7S821, D20S484, MFGT24, and MFGT27) in 10 μl reaction volumes comprising 2 μl of extracted fecal DNA and 5 μl of QIAGEN Multiplex PCR Master Mix (Multiplex PCR kit, Qiagen, USA) with 200–300 nM of each forward labeled and reverse non-labeled primer (set A, B, or C). The amplification parameters were as follows: 94 °C for 5 min; 45 cycles of 94 °C for 30 s, 57 °C for 30 s, and 60 °C for 30 min.

Amplification products were separated by capillary electrophoresis using an ABI 3130xl Genetic Analyzer (Applied Biosystems, CA, USA). Alleles were sized using Peak Scanner (Applied Biosystems). Since DNA extracted from non-invasively collected samples is typically degraded and low in concentration (Tarberlet et al. 1996; Morin et al. 2001; Ishizuka et al. 2019), we repeated the genotyping to ensure accuracy. According to Arandjelovic et al. (2009), the number of PCR replicates needed was twice when the allelic dropout rate was 5.2%. Since the allelic dropout rate at every locus was under 5% (see Table S2), we adopted the following criteria: homozygotes when a single allele was repeatedly observed at least three times at each locus or heterozygotes when each allele was repeatedly observed more than twice at each locus. Genotype accuracy and sample identification were confirmed by verifying that all mother–offspring pairs shared an allele at each locus or using two independently collected samples per individual. Although only one sample was successfully collected for two males, we included their genotype data because we directly observed their defecation and their genotypes were different from others.

### Paternity analysis

The paternity of 46 offspring (8 in 2018, 9 in 2019, 8 in 2020, 10 in 2021, and 11 in 2022 respectively) was investigated. Paternity analysis was conducted using the pairwise likelihood approach with CERVUS (Marshall et al. 1998). The resident candidate fathers were only fully sampled for the offspring born in 2020, 2021, and 2022, thus the percentage of EGP can be calculated for only these years. According to our field observations, there seemed to be approximately10 unsampled candidate fathers who belonged to the group during the 2017 and/or 2018 mating seasons, or were non-resident males. Since we collected genetic samples for 19 candidate fathers, which was 60–70% of all candidate fathers, the proportion of sampled candidate fathers was eventually assumed to be 0.60. The proportion of loci mistyped and the error rate in the likelihood calculation was both 0.01 because genotype accuracy was confirmed following the reasonable criteria (Arandjelovic et al. 2009). Paternity for 42 of the 46 offspring was analyzed under the condition that the offspring’s maternity was known since genotypes for the mothers were determined. However, genetic samples for mothers of four offspring were unsuccessfully collected. Therefore, the paternity for the four offspring was analyzed under the condition that the maternity of the offspring were unknown. When the most likely father had no mismatched alleles and the confidence level for the assignments was more than 95%, the male was concluded to be the offspring’s father.

### Statistical analysis

To analyze the likelihood of siring by each male, we constructed generalized linear mixed models (GLMMs) with a binomial error structure. We included the percentage of sires for each male in each year as a response variable (n=57). The social status (dominant/subordinate/non-resident) and age class (adult/subadult) of each male and the number of adult females at each year (in 2017, the number of adult females present during the mating season was an approximate value due to incomplete individual identification) were included as predictor variables. Offspring ID and the study year were included as random effects. We calculated the Akaike’s information criterion (AIC) for the constructed model and null model. Comparing the AIC values, we assessed the fit of constructed model. The models were fitted using the “lmer” function of the R package lme4 (Bates et al. 2015).

## Results

### Paternity

The genotypes of 87 individuals were analyzed. The mean proportion of loci typed was 1.00. The mean observed error rate was 0.018. The mean expected heterozygosity was 0.66 (see Table S2 in detail). The paternity of 34 of the 46 offspring was assigned to a single candidate father (Figure 1 and Table S3). When offspring born between 2019 and 2021 were conceived, “SN” was the alpha male. The percentage of his sires gradually decreased from 33% (3 of 9) in 2019, 13% (1 of 8) in 2020 to 0% (0 of 10) in 2021. When offspring born in 2022 were conceived, “TY” was the alpha male, and he sired 9% (1 of 11) of the offspring. The overall percentage of alpha male paternity was 13% (5 of 38). The alpha male at the time when the offspring of 2018 were born was not defined due to the complex changes in the alpha status (see the Materials and Methods).

**Fig. 1.**
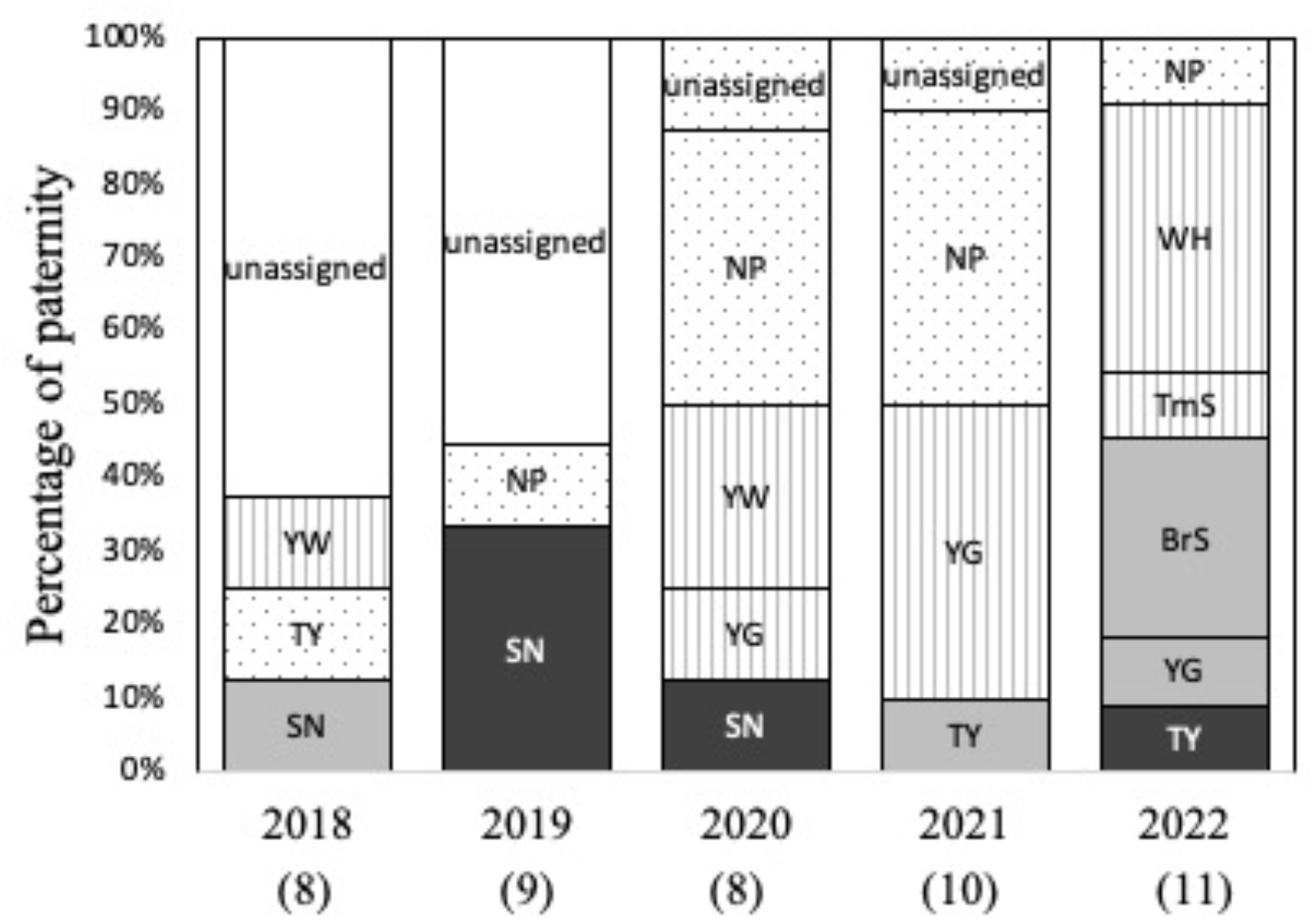
The number of offspring sired by each male in the five-year study period. Black, gray, lined, and dot bars indicate sires by the alpha, dominant, subordinate, and non-resident males, respectively. White bars indicate sires whose father’s social status remained unclear. The ID of all fathers is shown in each bar.

At least one offspring was sired by sampled non-resident males in each year. Since all resident candidate fathers for offspring born in 2018 and 2019 were not sampled, it remains unclear whether or not these offspring were sired by non-resident males, and thus, the percentage of EGP could not be calculated for those years. However, for the years in which all resident candidate fathers were sampled reveal that the percentage of EGP was 50% (4 of 8) in 2020, 50% (5 of 10) in 2021, and 9% (1 of 11) in 2022. The overall percentage of EGP between 2020 and 2022 was 34% (10 of 29). Interestingly, 75% (3 of 4) and 80% (4 of 5) of the EGP cases were assigned to a single non-resident male “NP” in 2020, 2021, respectively. He was the most successful sire and sired 33% (3 of 9) and 40% (4 of 10) of offspring in 2020 and 2021, respectively.

### Effects of male categories on the percentages of paternity success

The constructed model was significantly better fitted than the null model (ΔAIC = 10.38, P < 0.01). The constructed model showed that the effect of male age class had the negative effect on the percentage of paternity success, indicating that subadult males gain paternity success less likely compared to the adult males (Figure 2 and Table 1). The effect of male social status was not significant (Table 1).

**Table 1.** Results of the generalized linear mixed model (GLMM) performed to test the effects of males’ social status with their age class

| Parameter | Log-Odds | SE | 95% CI | Z | P |
| --- | --- | --- | --- | --- | --- |
| (Intercept) | -1.53 | 1.15 | -3.78, 0.73 | -1.33 | 0.184 |
| Male category (non-resident<br>vs. dominant) | -0.45 | 1.07 | -2.55, 1.65 | -0.42 | 0.677 |
| Male category (non-resident<br>vs. subordinate) | -0.36 | 1.06 | -2.44, 1.71 | -0.34 | 0.733 |
| Age (adult vs. subadult) | -2.97 | 1.04 | -5.00, -0.93 | -2.86 | < 0.01 |
| No. of adult females | -0.03 | 0.03 | -0.10, 0.03 | -1.04 | 0.300 |

**Fig. 2.**
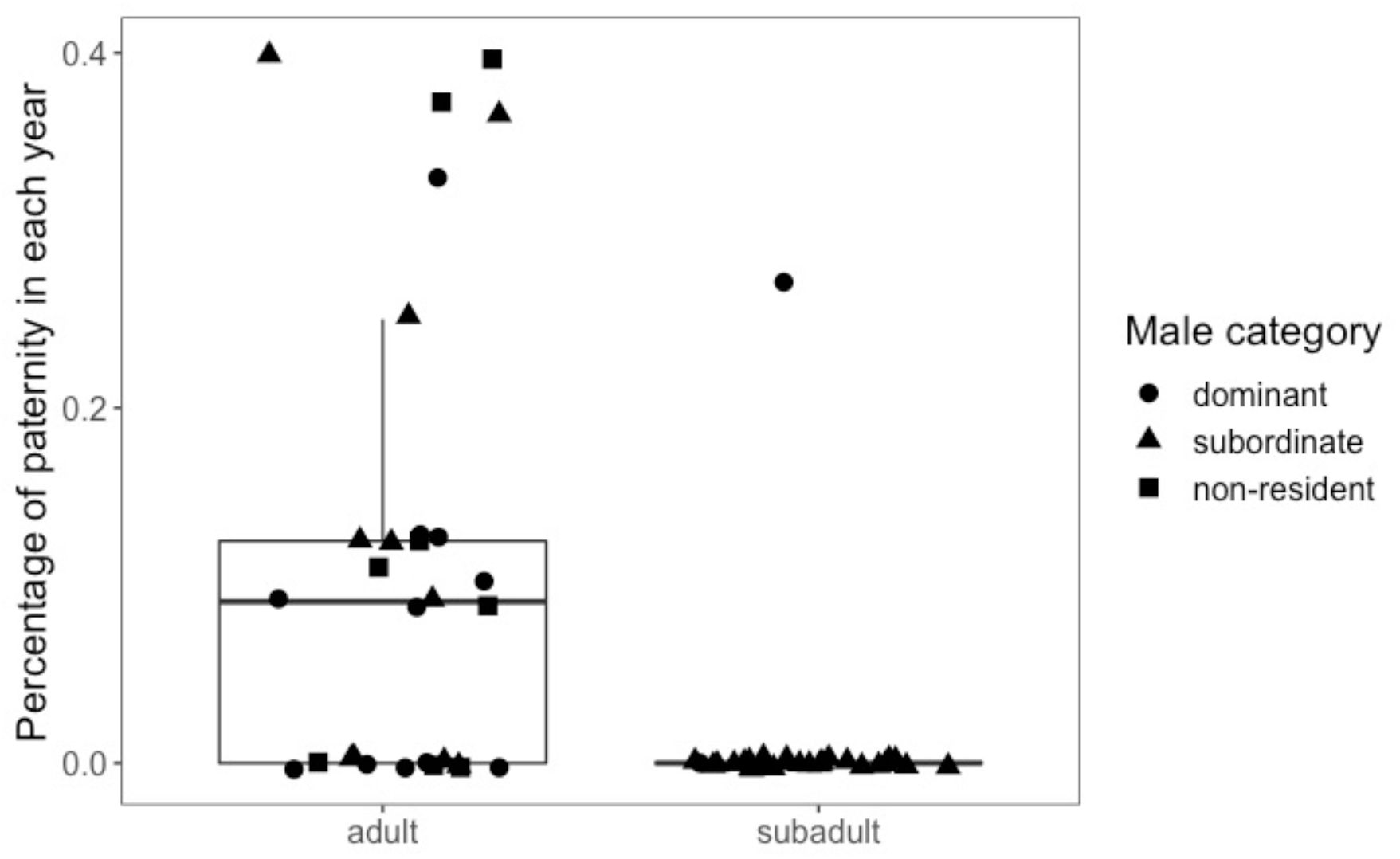
Percentage of paternity success by each male in each year with regards to the males’ social status. Boxes indicate the first to third quartile of observed values, solid lines show the median, and each dot represents the percentage of paternity success by a sampled candidate father in each year.

### The proportion of paternal sibling dyads

The percentages of paternal sibling dyads among the offspring born in 2018, 2019, 2020, 2021, and 2022 were 0%, 8%, 14%, 27%, and 16%, respectively (Table 2). The overall percentage of paternal sibling dyads was 15%. The overall proportion of paternal sibling dyads produced by the alpha male was only 2% and 0% in four of the five study years. Interestingly, one of subordinate males or non-resident males produced 11–13% of paternal sibling dyads between 2020 and 2022.

**Table 2.** The percentage of paternal sibling dyads among offspring born in each year. The values within the brackets represent the value of paternal sibling dyads in each category.

| Year | No. of dyads analyzed | Percentage of paternal sibling dyads | Percentage of paternal sibling dyads produced by the alpha male | Maximum percentage of paternal sibling dyads produced by one male | Male category of the most successful sire |
| --- | --- | --- | --- | --- | --- |
| 2018 | 28 | 0(0)* | 0 (0) | N/A | At least not alpha |
| 2019 | 36 | 8(3)* | 8 (3) | N/A | N/A |
| 2020 | 28 | 14(4) | 0 (0) | 11(3) | NR |
| 2021 | 45 | 27(12) | 0(0) | 13(6) | S and NR |
| 2022 | 55 | 16(9) | 0(0) | 11(6) | S |
| Total | 192 | 15(28) | 2 (3) | 12(15) | - |

S, and NR represents subordinate, and non-resident males, respectively. N/A represents data that was not available.
\* The percentages may be underestimated because offspring whose fathers remain unidentified may be paternal siblings with each other.

## Discussion

The percentage of EGP between 2020 and 2022 was 34% (9–50%) in the Shodoshima B-group. These results are consistent with previous results showing that 23–80% of offspring were sired by non-resident males (Table 3). The percentage of EGP in this species is relatively high among mammalian species (Isvaran and Clutton-Brock 2007; Soulsbury 2010). One reason for the relatively high percentage of EGP may be the high extent of female receptive synchrony in this species (Soltis et al. 2001). EGP is more likely when the number of males within groups decreases in mammals (Isvaran and Clutton-Brock 2007; Ostner et al. 2008; Ruiz-Lambides et al. 2017) and when female reproductive synchrony increases (Isvaran and Clutton-Brock 2007; Ostner et al. 2008; Ruiz-Lambides et al. 2017). Similar to other species that exhibit both high female receptive synchrony and EGP, resident males may fail to guard receptive females against non-resident males.

**Table 3.**
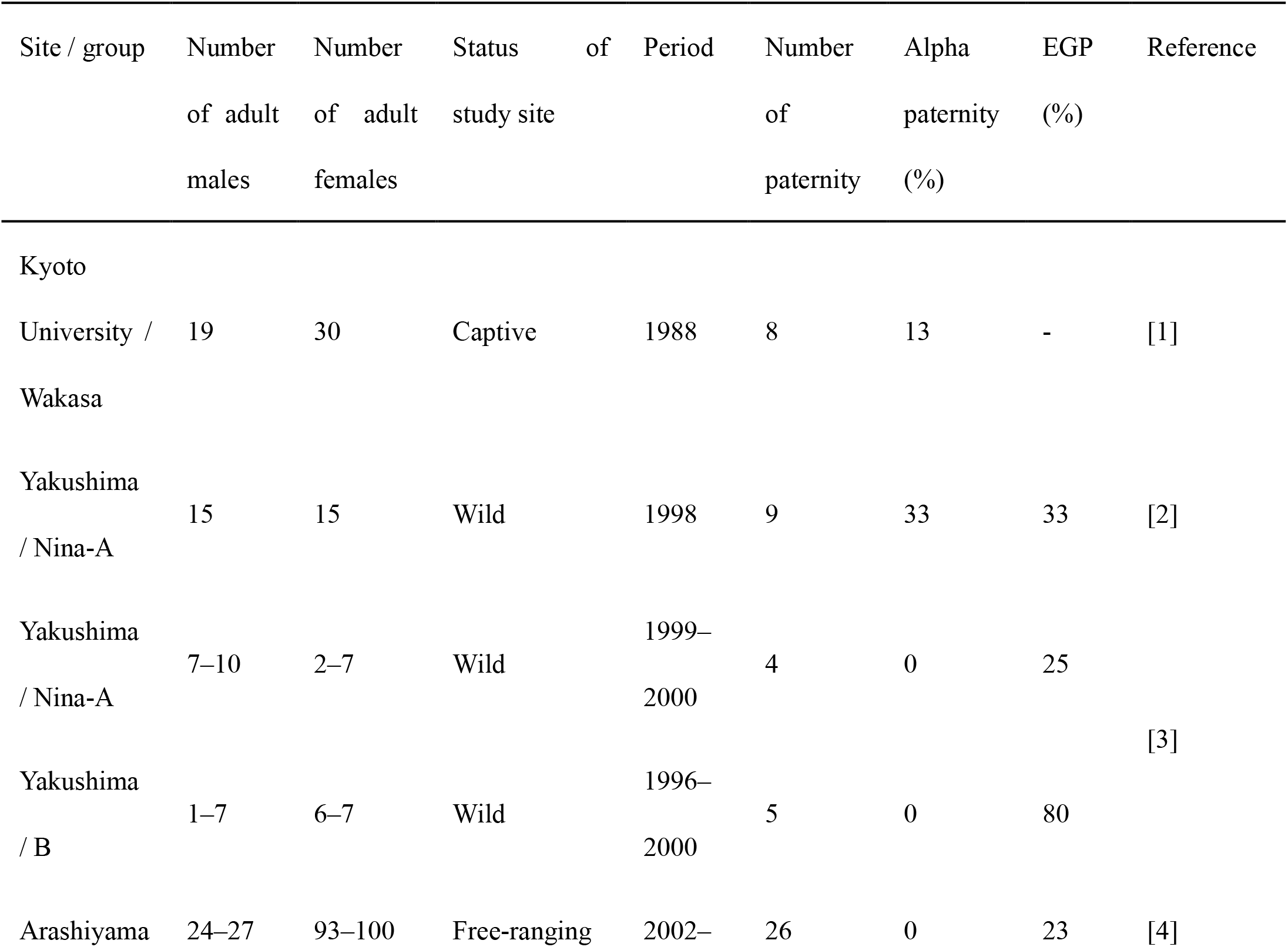
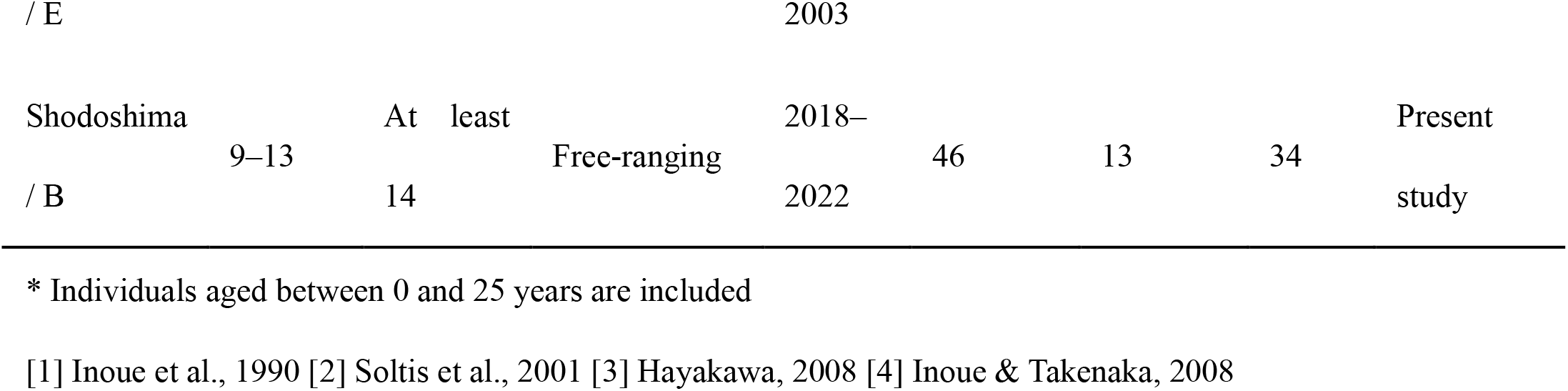
Overview of paternity results in Japanese macaques

An non-resident male, “NP,” was the most successful sire in both 2020 and 2021. Our results highlighted that non-resident males did not have a lower success rate with respect to siring offspring compared to resident males (including both dominant and subordinate males). These results clearly show that non-resident males have chances comparable to that of resident males when it comes to gain paternity success. Given that non-resident males might have chances to breed with females of other groups, the percentage of paternity success for non-resident males in this study might be underestimated. Since several social mammals prefer to mate with unfamiliar males (e.g. rodents: Kozakiewicz et al. 2009; primates: Huffman 1987), non-resident males may maintain unfamiliarity with females for at least several years, and consequently have several chances to breed with females. Furthermore, our results newly suggest that there is a reproductive skew among non-resident males; 75–80% of EGP were attained by “NP” in 2020 and 2021, while “TN” was unsuccessful in siring offspring in those two years. According to our field observations, “NP” was a large prime-aged male that seemed relatively dominant over other non-resident males, including “TN.” Since non-resident males are typically unfamiliar to females, it may not be female choice that triggers the reproductive skew among non-resident males. Instead, as the more dominant males often gain higher paternity success in various group-living mammalian species (e.g. ungulates: Pemberton et al. 1992; carnivores: Girman et al. 1997; primates: Ishizuka et al. 2018), reproductive skew among non-resident males may be mediated by dominance hierarchies among themselves.

The percentage of alpha male paternity was 13% (0–33%) in the Shodoshima-B group. These results were consistent with previous results where the percentages of alpha male paternity was 0–33% in this species (Table 3). Compared to other mammalian species, the percentage of the alpha male paternity is relatively low in Japanese macaques (Ostner et al. 2008; Gogarten and Koenig 2013; Soulsbury 2010), which suggests that male reproductive success is not highly skewed towards alpha males in this species. One possible reason for this is due to female mate choice. Females of this species are known to be highly selective when it comes to mating partners for breeding (Soltis et al. 1999; Inoue and Takenaka, 2008; Gartland et al. 2021). In our results, the percentage of offspring sired by “SN,” the alpha male during the mating seasons in 2018, 2019, and 2020, gradually decreased as the years passed (33% in 2019, 13% in 2020, 0% in 2021). This could be because he might have become more familiar and therefore avoided by females for mating as his alpha status tenure becomes longer. Female choice has been shown to negatively impact alpha males’ siring success in mammalian species, such as bank voles *Myodes glareolus* (Kozakiewicz et al. 2009). For females, unfamiliar males tend to be genetically dissimilar males; therefore, females may prefer to mate with unfamiliar males to gain genetic benefits for their offspring (Potts et al. 1991; Mays and Hills 2004).

Notably, subadult males sired a lower percentage of offspring than the adult males. These results were consistent with previous studies that report that young males that were physically immature sired only a few offspring in macaque species (Bercovitch et al. 2003; Inoue and Takenaka 2008; Dubuc et al. 2014a). Previous studies on rhesus macaques suggest that the darker facial coloration of adult males is sexually more attractive to females (Waitt et al. 2003; Dubuc et al. 2014b; Petersdorf et al. 2017). In Japanese macaques, the facial coloration of males appears to become darker as they mature, suggesting that females might show a similar preference to adult-aged males with dark facial coloration.

The percentage of paternal sibling dyads among offspring in the same age cohort was 15% (8–27%) in the Shodoshima-B group. Although available data for comparisons with our results is scarce, the percentage is relatively high compared to the 12% in rhesus macaques (Widdig 2013) and 5% in Assamese macaques *Macaca assamensis* (De Moor et al. 2020). These are noteworthy given that the percentage of alpha male paternity obtained in this study was lower (13%) than in rhesus macaques (24%, Widdig et al. 2004) and Assamese macaques (29%, Sukmak et al. 2014). In fact, alpha males produced no paternal sibling dyads in the same age cohorts in four of the five years of observation, and the percentage of paternal sibling dyads in the same age cohort produced by alpha males in the other year was only 2%. These results suggest that various males, including non-resident males, contribute more significantly to the production of paternal sibling dyads in Japanese macaques. The presence of paternal sibling dyads in the same age cohort may trigger kin selection and enhance affinity among individuals of similar age. In Japanese macaques, newly innovated behaviors are more easily transmitted among similar-aged individuals than among different-aged individuals (Kawai 1965; Huffman 1984), suggesting that paternal siblings in cohorts of the same age are affiliated with each other. Such an affinity among similar-aged individuals in Japanese macaques may be supported by the relatively high percentage of paternal sibling dyads in the same age cohorts. Future behavioral and genetic studies are required to clarify the presence or absence of paternal kin bias in this species.

Our findings contribute to a better understanding of the diverse reproductive strategies of males concerning their social status. Notably, this study clearly showed that the non-resident males have chances comparable to that of resident males when it comes to gaining paternity success. Furthermore, various males, including non-resident males, significantly contribute to the production of paternal sibling dyads in the same age cohorts, suggesting that not only the resident males, but also non-resident males play an important role in shaping within-group kin structures. However, future studies are required to examine how paternal siblings interact with each other.

## Statements and Declarations

### Data availability

The data that support the results of the study are available in the Supplementary Information.

### Author contributions

SI designed this study, collected behavioral and genetic data, analyzed all data, and wrote the manuscript. EI supported to design this study and write the manuscript. All authors gave final approval for publication.

### Ethical note

This study was permitted by the Choshikei Monkey Park on Shodoshima Island, Japan. All methods used in this study were noninvasive. Field research was conducted in accordance with the American Society of Primatologists Code of Best Practices for Field Primatology, and conformed to the Guidelines for Field Research established by the Ethics Committee of the Primate Research Institute of Kyoto University. All aspects of this study adhered to the ASAB/ABS Guidelines for the use of animals in research.

## Acknowledgements

We thank Dr. K. Watanabe, Mr. A. Nishio, Ms. C. Saeki, Mr. M. Ishii, Ms. Y. Kaji, Ms. K. Miyashita, and Mr. K. Hida for their help with the fieldwork. This study was financially supported by the Japan Society for the Promotion of Science Grant-in-Aid for JSPS fellows (21J00922 to SI), Cooperative Research Program of the Wildlife Research Center, Kyoto University (2020-B-05 to SI), and Leading Graduate Program in Primatology and Wildlife Science of Kyoto University.

## References

1. Altmann J (1979) Age cohorts as paternal sibships. Behav Ecol Sociobiol 6:161–164. https://doi.org/10.1007/BF00292563

2. Arandjelovic M, Guschanski K, Schubert G, Harris TR, Thalmann O, Siedel H, Vigilant L (2009) Two-step multiplex polymerase chain reaction improves the speed and accuracy of genotyping using DNA from noninvasive and museum samples. Mol Ecol 9:28–36 http://dx.doi.org/10.1111/j.1755-0998.2008.02387.x

3. Bates D, Maechler M, Bolker B, Walker S (2015) Fitting linear mixed-effects models using lme4. J Stat Softw 67:1–48

4. Bercovitch FB, Widdig A, Trefilov A, Kessler MJ, Berard JD, Schmidtke J, Nürnberg P, Krawczak M (2003) A longitudinal study of age-specific reproductive output and body condition among male rhesus macaques, Macaca mulatta. Naturwissenschaften 90:309–312. https://doi.org/10.1007/s00114-003-0436-1

5. Cords M, Minich T, Roberts SJ, Sleator C (2018) Evidence for paternal kin bias in the social affiliation of adult female blue monkeys. Am J Primatol 80:e22761. https://doi.org/10.1002/ajp.22761

6. Cowlishaw G, Dunbar RIM (1991) Dominance rank and mating success in male primates. Anim Behav 41:1045–1056. https://doi.org/10.1016/S0003-3472(05)80642-6

7. Dubuc C, Ruiz-Lambides A, Widdig A (2014a) Variance in male lifetime reproductive success and estimation of the degree of polygyny in a primate. Behav Ecol 25:878–889. https://doi.org/10.1093/beheco/aru052

8. Dubuc C, Allen WL, Maestripieri D, Higham JP (2014b) Is male rhesus macaque red color ornamentation attractive to females?. Behav Ecol Sociobiol 68:1215–1224. https://doi.org/10.1007/s00265-014-1732-9

9. Gartland KN, Biggs N, Shreeve CM, White FJ (2021) Dominance rank, female choice, and reproductive success in semi-free ranging adult male Japanese macaques (Macaca fuscata). Am J Primatol 83:e23294 https://doi.org/10.1002/ajp.23294

10. Girman DJ, Mills MGL, Geffen E, Wayne RK (1997) A molecular genetic analysis of social structure, dispersal, and interpack relationships of the African wild dog (Lycaon pictus). Behav Ecol Sociobiol 40:187–198. https://doi.org/10.1007/s002650050332

11. Gogarten JF, Koenig A (2013) Reproductive seasonality is a poor predictor of receptive synchrony and male reproductive skew among nonhuman primates. Behav Ecol Sociobiol 67:123–134. https://doi.org/10.1007/s00265-012-1432-2

12. Hamilton WD (1964) The genetical evolution of social behavior. J Theor Biol 7:1–52. https://doi.org/10.1016/0022-5193(64)90039-6

13. Hayakawa S (2008) Male–female mating tactics and paternity of wild Japanese macaques (Macaca fuscata yakui). Am J Primatol 70:986–989. https://doi.org/10.1002/ajp.20580

14. Hill DA (1999) Effects of provisioning on the social behaviour of Japanese and rhesus macaques: implications for socioecology. Primates 40:187–198. https://doi.org/10.1007/BF02557710

15. Huffman MA (1984) Stone-play of Macaca fuscata in Arashiyama B troop: transmission of a non-adaptive behavior. J Hum Evol 13:725–735. https://doi.org/10.1016/S0047-2484(84)80022-6

16. Huffman MA (1987) Consort intrusion and female mate choice in Japanese macaques (Macaca fuscata). Ethology 75:221–234. https://doi.org/10.1111/j.1439-0310.1987.tb00655.x

17. Inoue E, Takenaka O (2008) The effect of male tenure and female mate choice on paternity in free-ranging Japanese macaques. Am J Primatol 70:62–68. https://doi.org/10.1002/ajp.20457

18. Inoue M, Takenaka A, Tanaka S, Kominami R, Takenaka O (1990) Paternity discrimination in a Japanese macaque group by DNA fingerprinting. Primates 31:563–570. https://doi.org/10.1007/BF02382539

19. Ishizuka S, Kawamoto Y, Sakamaki T, Tokuyama N, Toda K, Okamura H, Furuichi T (2018) Paternity and kin structure among neighbouring groups in wild bonobos at Wamba. R Soc Open Sci 5:171006. https://doi.org/10.1098/rsos.171006

20. Ishizuka S, Kawamoto Y, Toda K, Furuichi T (2019) Bonobos’ saliva remaining on the pith of terrestrial herbaceous vegetation can serve as non-invasive wild genetic resources. Primates 60:7–13. https://doi.org/10.1007/s10329-018-00704-x

21. Ishizuka S (2021) Do dominant monkeys gain more warmth? Number of physical contacts and spatial positions in huddles for male Japanese macaques in relation to dominance rank. Behav Process 185:104317 https://doi.org/10.1016/j.beproc.2021.104317

22. Isvaran K, Clutton-Brock T (2007) Ecological correlates of extra-group paternity in mammals. Proc R Soc Lond B 274:219–224. https://doi.org/10.1098/rspb.2006.3723

23. Kawai M (1965) Newly acquired pre-cultural behavior of the natural troop of Japanese monkeys on Koshima Islet. Primates 6:1–30.

24. Kawazoe T (2016) Association patterns and affiliative relationships outside a troop in wild male Japanese macaques, Macaca fuscata, during the non-mating season. Behaviour 153:69–89. https://doi.org/10.1163/1568539X-00003325

25. Kozakiewicz M, Choŀuj A, Kozakiewicz A, Sokóŀ M (2009) Familiarity and female choice in the bank vole—do females prefer strangers?. Acta Theriol 54:157–164. https://doi.org/10.1007/BF03193171

26. Kutsukake N, Nunn CL (2009) The causes and consequences of reproductive skew in male primates. In: Hager R, Jones CB (eds) Reproductive skew in vertebrates: proximate and ultimate causes. Cambridge University Press, Cambridge, pp 165–195

27. Langergraber KE, Mitani JC, Vigilant L (2007) The limited impact of kinship on cooperation in wild chimpanzees. Proc Natl Acad Sci USA 104:7786–7790. https://doi.org/10.1073/pnas.0611449104

28. Lukas D, Reynolds V, Boesch C, Vigilant L (2005) To what extent does living in a group mean living with kin?. Mol Ecol 14:2181–2196. https://doi.org/10.1111/j.1365-294X.2005.02560.x

29. Lyke MM, Dubach J, Briggs MB (2013) A molecular analysis of African lion (Panthera leo) mating structure and extra-group paternity in Etosha National Park. Mol Ecol 22:2787–2796. https://doi.org/10.1111/mec.12279

30. Marshall, T. C., Slate, J., Kruuk, L. E., & Pemberton, J. M. (1998). Statistical confidence for likelihood-based paternity inference in natural populations. Mol Ecol 7:639–655 https://doi.org/10.1046/j.1365-294x.1998.00374.x

31. Maruhashi T (1982) An ecological study of troop fissions of Japanese monkeys (Macaca fuscata yakui) on Yakushima Island, Japan. Primates 23:317–337. https://doi.org/10.1007/BF02381317

32. Mays Jr HL, Hill GE (2004) Choosing mates: good genes versus genes that are a good fit. Trends Ecol Evol 19:554–559. https://doi.org/10.1016/j.tree.2004.07.018

33. Miller CM, Snyder-Mackler N, Nguyen, N et al (2021) Extragroup paternity in gelada monkeys, Theropithecus gelada, at Guassa, Ethiopia and a comparison with other primates. Anim Behav 177:277–301. https://doi.org/10.1016/j.anbehav.2021.05.008

34. Morin PA, Chambers KE, Boesch C, Vigilant L (2001) Quantitative polymerase chain reaction analysis of DNA from noninvasive samples for accurate microsatellite genotyping of wild chimpanzees (Pan troglodytes verus). Mol Ecol 10:1835–1844 https://doi.org/10.1046/j.0962-1083.2001.01308.x

35. Nichols HJ, Cant MA, Sanderson JL (2015) Adjustment of costly extra-group paternity according to inbreeding risk in a cooperative mammal. Behav Ecol 26:1486–1494. https://doi.org/10.1093/beheco/arv095

36. Ortega J, Maldonado JE, Wilkinson GS, Arita HT, Fleischer RC (2003) Male dominance, paternity, and relatedness in the Jamaican fruit-eating bat (Artibeus jamaicensis). Mol Ecol 12: 2409–2415. https://doi.org/10.1046/j.1365-294X.2003.01924.x

37. Ostner J, Nunn CL, Schülke O (2008) Female reproductive synchrony predicts skewed paternity across primates. Behav Ecol 19:1150–1158 https://doi.org/10.1093/beheco/arn093

38. Pemberton JM, Albon SD, Guinness FE, Clutton-Brock TH, Dover GA (1992) Behavioral estimates of male mating success tested by DNA fingerprinting in a polygynous mammal. Behav Ecol 3:66–75. https://doi.org/10.1093/beheco/3.1.66

39. Petersdorf M, Dubuc C, Georgiev AV, Winters S, Higham JP (2017) Is male rhesus macaque facial coloration under intrasexual selection?. Behav Ecol 28:1472–1481. https://doi.org/10.1093/beheco/arx110

40. Potts WK, Manning CJ, Wakeland EK (1991) Mating patterns in seminatural populations of mice influenced by MHC genotype. Nature 352:619–621. https://doi.org/10.1038/352619a0

41. Reed DH, Frankham R (2003) Correlation between fitness and genetic diversity. Conser Biol 17:230–237. https://doi.org/10.1046/j.1523-1739.2003.01236.x

42. Ruiz-Lambides AV, Weiß BM, Kulik L, Stephens C, Mundry R, Widdig A (2017) Long-term analysis on the variance of extra-group paternities in rhesus macaques. Behav Ecol Sociobiol 71:67 https://doi.org/10.1007/s00265-017-2291-7

43. Silk JB (2009) Nepotistic cooperation in non-human primate groups. Phil Trans R Soc Lond B 364:3243–3254. https://doi.org/10.1098/rstb.2009.0118

44. Smith K, Alberts SC, Altmann J (2003) Wild female baboons bias their social behaviour towards paternal half-sisters. Proc R Soc Lond B 270:503–510. https://doi.org/10.1098/rspb.2002.2277

45. Smith JE (2014) Hamilton’s legacy: kinship, cooperation and social tolerance in mammalian groups. Anim Behav 92:291–304. https://doi.org/10.1016/j.anbehav.2014.02.029

46. Soltis J, Mitsunaga F, Shimizu K, Yanagihara Y, Nozaki M (1999) Female mating strategy in an enclosed group of Japanese macaques. Am J Primatol 47:263–278 https://doi.org/10.1002/(SICI)1098-2345(1999)47:4<263::AID-AJP1>3.0.CO;2-F

47. Soltis J, Thomsen R, Takenaka O (2001) The interaction of male and female reproductive strategies and paternity in wild Japanese macaques, Macaca fuscata. Anim Behav 62:485–494 https://doi.org/10.1006/anbe.2001.1774

48. Soulsbury CD (2010) Genetic patterns of paternity and testes size in mammals. PLoS One 5:e9581. https://doi.org/10.1371/journal.pone.0009581

49. South JM, Emmons LH, Bernard H (2007) Behavioral monogamy and fruit availability in the large treeshrew (Tupaia tana) in Sabah, Malaysia. J Mamm 88:1427–1438. https://doi.org/10.1007/s00265-007-0454-7

50. Sukmak M, Wajjwalku W, Ostner J, Schülke O (2014) Dominance rank, female reproductive synchrony, and male reproductive skew in wild Assamese macaques. Behav Ecol Sociobiol 68:1097–1108. https://doi.org/10.1007/s00265-014-1721-z

51. Taberlet P, Griffin S, Goossens B, Questiau S, Manceau V, Escaravage N, Waits LP, Bouvet J (1996) Reliable genotyping of samples with very low DNA quantities using PCR. Nucleic Acids Res 24:3189–3194 https://doi.org/10.1093/nar/24.16.3189

52. Takahashi H (2001) Influence of fluctuation in the operational sex ratio to mating of troop and non-troop male Japanese macaques for four years on Kinkazan Island, Japan. Primates 42:183–191. https://doi.org/10.1007/BF02629635

53. Takahata Y, Suzuki S, Agetsuma N, Okayasu N, Sugiura H, Takahashi H, Yamagiwa J, Izawa K, Furuichi T, Hill DA, Maruhashi T, Saito C, Sato S, Sprague DS (1998) Reproduction of wild Japanese macaque females of Yakushima and Kinkazan islands: a preliminary report. Primates 39:339–349 https://doi.org/10.1007/BF02573082

54. Takahata Y, Huffman MA, Suzuki S, Koyama N, Yamagiwa J (1999) Why dominants do not consistently attain high mating and reproductive success: a review of longitudinal Japanese macaque studies. Primates 40:143–158 https://doi.org/10.1007/BF02557707

55. Toyoda A, Maruhashi T, Kawamoto Y, Matsudaira K, Matsuda I, Malaivijitnond S (2022) Mating and reproductive success in free-ranging stump-tailed macaques: effectiveness of male–male coalition formation as a reproductive strategy. Front Ecol Evol 10:802012. https://doi.org/10.3389/fevo.2022.802012

56. Wahaj SA, Van Horn RC, Van Horn TL, Dreyer R, Hilgris R, Schwarz J, Holekamp KE (2004) Kin discrimination in the spotted hyena (Crocuta crocuta): nepotism among siblings. Behav Ecol Sociobiol 56:237–247. https://doi.org/10.1007/s00265-004-0783-8

57. Waitt C, Little AC, Wolfensohn S, Honess P, Brown AP, Buchanan-Smith HM, Perrett DI (2003) Evidence from rhesus macaques suggests that male coloration plays a role in female primate mate choice. Proc R Soc Lond B 270:S144–S146. https://doi.org/10.1098/rsbl.2003.0065

58. Watanabe K (1979) Alliance formation in a free-ranging troop of Japanese macaques. Primates 20:459–474. https://doi.org/10.1007/BF02373429

59. Wells DA, Cant MA, Thompson FJ, Marshall HH, Vitikainen EI, Hoffman JI, Nichols HJ (2021) Extra-group paternity varies with proxies of relatedness in a social mammal with high inbreeding risk. Behav Ecol 32:94–104. https://doi.org/10.1093/beheco/araa105

60. Widdig A, Nürnberg P, Krawczak M, Streich WJ, Bercovitch FB (2001) Paternal relatedness and age proximity regulate social relationships among adult female rhesus macaques. Proc Natl Acad Sci USA 98:13769–13773. https://doi.org/10.1073/pnas.241210198

61. Widdig A, Bercovitch FB, Jürgen Streich W, Sauermann U, Nürnberg P, Krawczak M (2004) A longitudinal analysis of reproductive skew in male rhesus macaques. Proc R Soc Lond B 271:819–826 https://doi.org/10.1098/rspb.2003.2666

62. Widdig A (2007) Paternal kin discrimination: the evidence and likely mechanisms. Biol Rev 82:319–334 https://doi.org/10.1111/j.1469-185X.2007.00011.x

63. Widdig A (2013) The impact of male reproductive skew on kin structure and sociality in multi-male groups. Evol Anthropol 22:239–250. https://doi.org/10.1002/evan.21366

64. Wikberg EC, Jack KM, Fedigan LM et al (2017) Inbreeding avoidance and female mate choice shape reproductive skew in capuchin monkeys (Cebus capucinus imitator). Mol Ecol 26:653–667. https://doi.org/10.1111/mec.13898

65. Yamagiwa J, Hill DA (1998) Intraspecific variation in the social organization of Japanese macaques: past and present scope of field studies in natural habitats. Primates 39:257–273 https://doi.org/10.1007/BF02573076

