## Supplementary Information for "Comparisons of paternity success for resident and non-resident males and their influences on paternal sibling cohorts in Japanese macaques on Shodoshima Island"

**Table S1** Information for candidate fathers

| ID | 2017 | 2018 | | 2019 | | 2020 | | 2021 | | 2022 |
|---|---|---|---|---|---|---|---|---|---|---|
| | m | nm | m | nm | m | nm | m | nm | m | nm |
| SN | D/A | **D/A** | **D/A** | **D/A** | **D/A** | **D/A** | **D/A** | **D/A** | *absent* | *absent* |
| TY | NR/A* | D/A | D/A | D/A | D/A | D/A | D/A | D/A | **D/A** | **D/A** |
| YG | S/SA | S/A | S/A | S/A | S/A | S/A | S/A | S/A | D/A | D/A |
| YW | S/A | S/A | S/A | S/A | S/A | S/A | *absent* | *absent* | *absent* | *absent* |
| ClS | S/SA | S/SA | S/SA | S/SA | S/SA | S/A | S/A | *absent* | *absent* | *absent* |
| TR | D/A | *absent* | *absent* | *absent* | *absent* | *absent* | *absent* | *absent* | *absent* | *absent* |
| SB | D/A | *absent* | *absent* | *absent* | *absent* | *absent* | *absent* | *absent* | *absent* | *absent* |
| TmS | *immature* | S/SA | S/SA | S/SA | S/SA | S/SA | S/SA | S/A | S/A | S/A |
| BkS | *immature* | S/SA | S/SA | S/SA | S/SA | S/SA | S/SA | S/A | S/A | S/A |

| | | | | | | | | | | |
|---|---|---|---|---|---|---|---|---|---|---|
| WH | N/A | N/A | N/A | S/SA | S/SA | S/SA | S/SA | S/A | S/A | S/A |
| NP | N/A | N/A | NR/A | *absent* | NR/A | *absent* | NR/A | *absent* | NR/A | *absent* |
| TN | N/A | N/A | N/A | *absent* | NR/A | *absent* | NR/A | *absent* | NR/A | *absent* |
| BrS | *immature* | *immature* | *immature* | *immature* | *immature* | S/SA | D/SA | D/SA | D/SA | D/SA |
| BL | N/A | N/A | N/A | S/SA | S/SA | S/SA | S/SA | S/SA | S/SA | S/SA |
| KR | *immature* | *immature* | *immature* | S/SA | S/SA | S/SA | S/SA | S/SA | S/SA | S/SA |
| RZ | N/A | N/A | N/A | S/SA | S/SA | S/SA | S/SA | S/SA | S/SA | S/SA |
| HP | N/A | N/A | N/A | S/SA | S/SA | S/SA | S/SA | *absent* | *absent* | *absent* |
| NmS | *immature* | *immature* | *immature* | *immature* | *immature* | S/SA | S/SA | S/SA | S/SA | S/SA |
| HsS | *immature* | *immature* | *immature* | *immature* | *immature* | *immature* | *immature* | S/SA | S/SA | S/SA |

D, S, and NR indicates dominant, subordinate, and non-resident males, respectively. A, and SA indicates adult and subadult males, respectively.

Bold indicates the alpha male.

“m,” and “nm” indicates mating seasons and non-mating seasons, respectively.

Shadow indicates the candidate fathers.

* Although the demographic data for TY was not collected in non-mating season 2017, we classified the male as a non-resident male because he was begun to be observed in the end of September and was not habituated to human.

**Table S2** Summary for population genetic parameters

| Locus | N | No. of alleles | Observed heterozygosity | Expected heterozygosity | Allelic dropout rate |
|---|---|---|---|---|---|
| MFGT22 | 87 | 8 | 0.716 | 0.787 | 0% |
| D5s820 | 87 | 6 | 0.670 | 0.683 | 0% |
| MFGT18 | 87 | 6 | 0.489 | 0.456 | 0% |
| D6s501 | 87 | 8 | 0.875 | 0.826 | 0% |
| D3s1768 | 87 | 5 | 0.648 | 0.683 | 1% |
| D17s1290 | 87 | 8 | 0.693 | 0.712 | 1% |
| MFGT21 | 87 | 7 | 0.261 | 0.335 | 3% |
| MFGT5 | 87 | 3 | 0.443 | 0.471 | 1% |
| D6s493 | 87 | 7 | 0.716 | 0.713 | 1% |
| D14s306 | 87 | 5 | 0.670 | 0.712 | 0% |
| D19s582 | 87 | 6 | 0.693 | 0.667 | 1% |
| D20s484 | 87 | 7 | 0.625 | 0.778 | 4% |
| MFGT27 | 87 | 6 | 0.670 | 0.729 | 0% |
| D7s821 | 87 | 7 | 0.693 | 0.701 | 0% |
| MFGT24 | 87 | 7 | 0.795 | 0.751 | 1% |
| D1s548 | 87 | 4 | 0.511 | 0.478 | 0% |

**Table S3** Results for paternity of offspring in the B-group

| Offspring | Year of birth | Mother | Assigned father | Delta score | Confidence level |
|---|---|---|---|---|---|
| Km18 | 2018 | Km | - | - | - |
| Bs18 | | Bs | - | - | - |
| Sf18 | | Sf | - | - | - |
| Ma18 | | Ma | TY | 14.48 | > 95% |
| Dk18 | | Dk | YW | 6.00 | > 95% |
| Po18 | | Po | - | - | - |
| Bv18* | | Bv | - | - | - |
| Bb18* | | Bb | SN | 3.88 | > 95% |
| Bs19 | 2019 | Bs | - | - | - |
| Vg19 | | Vg | - | - | - |
| Ok19 | | Ok | - | - | - |
| Bk19 | | Bk | SN | 8.05 | > 95% |
| Rw19 | | Rw | - | - | - |
| Pe19 | | Pe | SN | 10.03 | > 95% |
| Sk19 | | Sk | - | - | - |
| Bb19* | | Bb | SN | 3.85 | > 95% |

| | | | | | |
|---|---|---|---|---|---|
| Tr19* | | Tr | NP | 10.42 | > 95% |
| Br20 | | Br | NP | 14.00 | > 95% |
| Km20 | | Km | - | - | - |
| Am20 | | Am | YG | 14.27 | > 95% |
| Rs20 | | Rs | YW | 9.76 | > 95% |
| Bs20 | 2020 | Bs | SN | 10.78 | > 95% |
| Nm20 | | Nm | NP | 18.58 | > 95% |
| Hk20 | | Hk | YW | 9.54 | > 95% |
| Rw20 | | Rw | NP | 16.04 | > 95% |
| Br21 | | Br | YG | 12.58 | > 95% |
| Rs21 | | Rs | NP | 20.35 | > 95% |
| Mr21 | | Mr | YG | 11.43 | > 95% |
| Qp21 | | Qp | NP | 16.93 | > 95% |
| Qb21 | | Qb | NP | 19.50 | > 95% |
| Tg21 | 2021 | Tg | YG | 12.79 | > 95% |
| Bs21 | | Bs | TY | 13.15 | > 95% |
| Vg21 | | Vg | NP | 19.14 | > 95% |
| Sf21 | | Sf | YG | 7.73 | > 95% |
| Ok21 | | Ok | - | - | - |

| | | | | | |
|---|---|---|---|---|---|
| Br22 | | Br | WH | 13.89 | > 95% |
| Km22 | | Km | WH | 13.00 | > 95% |
| Am22 | | Am | WH | 16.73 | > 95% |
| Qp22 | | Qp | TY | 13.02 | > 95% |
| Qb22 | | Qb | WH | 10.84 | > 95% |
| Bs22 | 2022 | Bs | BrS | 8.99 | > 95% |
| Sf22 | | Sf | BrS | 5.60 | > 95% |
| Nm22 | | Nm | NP | 16.95 | > 95% |
| Hk22 | | Hk | BrS | 8.21 | > 95% |
| Mm22 | | Mm | YG | 7.86 | > 95% |
| Mt22 | | Mt | TmS | 7.88 | > 95% |

* Samples for mothers of the individuals were not collected, so paternity analysis was performed under the condition where genotypes for their mothers were unavailable.
